# Cortical Oscillatory Dysrhythmias in Visual Snow Syndrome: A MEG Study

**DOI:** 10.1101/2021.05.17.444460

**Authors:** Jenny L. Hepschke, Robert A Seymour, Wei He, Andrew Etchell, Paul F Sowman, Clare L Fraser

**Affiliations:** Save Sight Institute, Faculty of Health and Medicine, The University of Sydney, Sydney, NSW Australia; Department of Ophthalmology, Prince of Wales Hospital, High Street, Randwick, NSW, Australia; Department of Cognitive Science, Macquarie University, Sydney NSW Australia; Macquarie Ophthalmology, Macquarie University, Sydney NSW; Wellcome Centre for Human Neuroimaging, UCL Queen Square Institute of Neurology, University College London, London WC1N 3AR, U.K

**Author notes:** Joint first authors (Jenny L. Hepschke, Robert A Seymour). **Corresponding Author:** Associate Professor Clare L Fraser, Save Sight Institute, 8 Macquarie Street, Sydney, NSW Australia.

**Keywords:** Visual Snow, Migraine, Dysrhythmia, Magnetoencephalography, Phase Amplitude Coupling

## Abstract

Visual Snow (VS) refers to the persistent visual experience of static in the whole visual field of both eyes. It is often reported by patients with migraine and co-occurs with conditions like tinnitus and tremor. The underlying pathophysiology of the condition is poorly understood. Previously we hypothesised, that VSS may be characterised by disruptions to rhythmical activity within the visual system^1^.

To test this, data from 18 patients diagnosed with visual snow syndrome (VSS), and 16 matched controls, were acquired using Magnetoencephalography (MEG). Participants were presented with visual grating stimuli, known to elicit decreases in alpha-band (8-13Hz) power and increases in gamma-band power (40-70Hz).

Data were mapped to source-space using a beamformer. Across both groups, decreased alpha power and increased gamma power localised to early visual cortex. Data from primary visual cortex (V1) were compared between groups. No differences were found in either alpha or gamma peak frequency or the magnitude of alpha power, p>.05. However, compared with controls, our VSS cohort displayed significantly increased V1 gamma power, p=.035. This new electromagnetic finding concurs with previous fMRI and PET findings suggesting that in VSS, the visual cortex is hyper-excitable. The coupling of alpha-phase to gamma amplitude (i.e., phase-amplitude coupling, PAC) within V1 was also quantified. Compared with controls, the VSS group had significantly reduced alpha-gamma PAC, p<.05, indicating a potential excitation-inhibition imbalance in VSS, as well as a potential disruption to top-down “noise-cancellation” mechanisms.

Overall, these results suggest that rhythmical brain activity in primary visual cortex is both hyperexcitable and disorganised in VSS, consistent with visual snow being a condition of thalamocortical dysrhythmia.

## Introduction

Visual Snow (VS) refers to the persistent visual experience of static in the whole visual field of both eyes, likened to “static analogue television noise”.^2^ This phenomena was initially reported by patients with migraine^3^ but more recently has been classified as a syndrome with specific diagnostic criteria to capture the spectrum of the pathology.^4,5^ Visual snow syndrome (VSS) is defined as flickering fine achromatic dots with at least one associated visual symptom of palinopsia, photopsia, nyctalopia, and entoptic phenomena, as well as non-visual symptoms such as tinnitus, migraine, and tremor. ^4,6^ Previous epidemiological studies have shown that VSS exists as a continuum and that the frequency of associated non-visual symptoms often carries a higher symptom severity and burden of disease.^6–8^ The condition has an estimated prevalence of around 2% in the United Kingdom.^9,10^

To date, the pathophysiology underlying VSS is poorly understood, though the high co-prevalence of migraine and tinnitus suggests it may be a disorder of sensory processing.^1,10^ In support of this, recent neuroscientific work has demonstrated various functional and structural alterations within the primary visual cortex (V1),^6^ and ventral visual regions,^11^ of VSS patients. Co-occurring hypermetabolism and cortical volume increases at the intersection of right lingual and fusiform gyrus have also recently been reported.^8^ Resting-state functional MRI data from a VSS cohort showed hyperconnectivity between extrastriate and inferior temporal brain regions and prefrontal and parietal regions.^12^ VSS patients also demonstrate variations in visual evoked potentials,^13^ as well as disrupted habituation for repeated stimuli.^14^ Overall, there is an emerging picture of co-occurring visual hyperactivity, hyperconnectivity, and dishabituation in VSS that could result from a faulty “noise-cancelling” mechanism,^15^ similar to that in the auditory domain for tinnitus.^16,17^

Our group has recently proposed that VSS symptoms may be underpinned by perturbations to the rhythms of the human visual system,^1^: in particular, a disruption to the usual, state-dependent, flow of information within the thalamocortical network. Successful perceptual processing relies upon the coordinated activity of large groups of neuronal cell assemblies throughout the brain, firing in a rhythmic fashion.^18–20^ These neuronal “oscillations” can be measured outside the head non-invasively using EEG or MEG.^21^ We hypothesise that disruptions to visual oscillations may represent a central pathophysiological mechanism in VSS. Specifically, visual dysrhythmia could alter cortical circuit entrainment and top-down control in VSS, thereby altering the threshold for transmission, affecting suppression and attention, and allowing for detection of sub-threshold visual stimuli.^22,23^ Similar disruptions to the endogenous sensory rhythms of the brain are found in other conditions associated with sensory defects, including migraine, neuropathic pain, and tinnitus.^24–26^

This study aimed to investigate the dysrhythmia hypothesis by studying endogenous rhythmical activity (neural oscillations) in the visual system of VSS patients versus controls. We focussed on oscillations in two frequency bands. First, gamma-band (40-100Hz) oscillations; generated locally via the coordinated interaction between excitatory and inhibitory populations of neurons.^27^ These oscillations are thought to provide a precise timing mechanism,^28^ to facilitate information transfer up the cortical hierarchy.^29^ Alterations in gamma-band activity have been reported for other conditions of ‘phantom’ perception, including tinnitus,^30–32^ and neuropathic pain.^33^ Given the reports of hyperexcitability in VSS,^8^ we expected patients to show increased gamma-band power. The second frequency band of interest was the alpha band (8-13Hz). Alpha rhythms are widely observed in EEG and MEG recordings, originating from several cortical and thalamic generators.^20,34^ Alpha power is negatively correlated with sustained attention and is involved in the active inhibition of irrelevant visual information.^35^ There is emerging evidence that alpha-band oscillations are also involved in long-range functional connectivity,^20^ and the modulation of local gamma oscillations within the visual cortex via a phase-amplitude coupling.^36,37^ Given the hypothesised reduction in a top-down, ‘noise cancellation’ mechanism,^1,25^ we expected VSS patients to show reductions in the modulation of local gamma oscillations via alpha-band phase.

We tested these hypotheses using magnetoencephalography (MEG) combined with a simple visual-grating paradigm known to elicit reliable changes in both alpha and gamma oscillations in the primary visual cortex.

## Materials and Methods

### Participants

Eighteen patients with Visual Snow Syndrome (VSS) and 16 age- and gender-matched controls participated in this study. Before MEG, potential VSS patients underwent a comprehensive neuro-ophthalmic examination including a standardised series of questions about visual and non-visual symptoms to establish VSS duration, associated features and previous diagnoses. This included the measurement of visually evoked potential (VEP), pattern electroretinogram (pERG) and fullfield Electroretinogram (ffERG). VSS participants were included if they fulfilled the diagnostic criteria of typical VSS.^4^ Participants were excluded if they were taking psychiatric medication, reported epileptic symptoms, had a diagnosis of hallucinogen-persistence perceptual disorder (HPPD), showed any abnormality on brain MRI or visual electrophysiology.

### Experimental Procedures

Experimental procedures complied with the Declaration of Helsinki and were approved by Macquarie University Human Research Ethics Committee. Written consent was obtained from all participants.

### Experimental Paradigm and Design

Participants performed a visual task (Figure 1) while their brain activity was continuously recorded with MEG. The task contained an embedded black and white visual grating stimulus that has been shown to reliably elicit gamma-band oscillations.^38,39^ Each task trial started with a fixation period (2.0, 3.0, or 4.0s), followed by a monochrome visual grating (spatial frequency of 2 cycles/degree) for 1.5s. Following this, a cartoon picture of an alien or astronaut was presented for 1.0s. This segment of the trial was included only to maintain the engagement and arousal of the participant; the neural response to this stimulus was not analysed. At the end of the trial, participants were presented with a question mark (‘?’) and instructed to respond if they had just seen an alien picture using a response pad (maximum response period of 1.0 s). Feedback about the correctness of responses was conveyed to the participant via a short (0.1s) auditory tone. MEG recordings lasted 15-16 minutes and included 150 trials. Accuracy rates were >95% for all participants.

**Figure 1:**
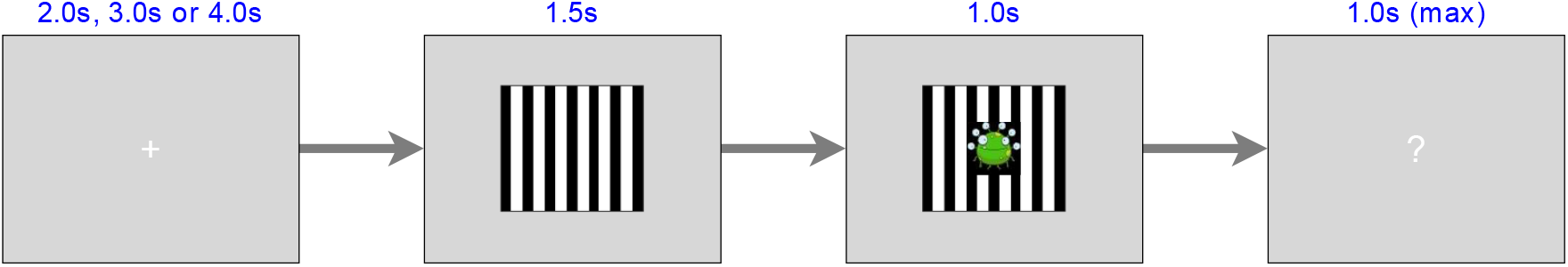
Experimental paradigm. Following a 2.0s, 3.0s, or 4.0s baseline period, participants were presented with a visual grating (1.5s duration). A cartoon alien or astronaut picture (duration 1.0s) was then presented. The subsequent presentation of a ‘?’ symbol was the imperative signal for a response to an alien (response time up to 1.0 s). Participants were instructed to provide no response to astronauts. The alien/astronaut stimuli were to maintain attention and were not part of the analysed data.

### MEG Acquisition

Data were acquired using a KIT MEG160 magnetoencephalograph (Model PQ1160R-N2, KIT, Kanazawa, Japan) consisting of 160 coaxial first-order gradiometers with a 50mm baseline. The KIT MEG160 is arranged in a fixed supine acquisition configuration and is located within a magnetically shielded room (Fujihara Co. Ltd., Tokyo, Japan). Continuous MEG, within a passband of 0.03–200 Hz, was sampled at 1000 Hz. Five head position indicator or “marker” coils were applied for head position measurement, and measurements were taken from these before and after the experiment. No participant moved more than 5mm in any direction (x, y, z) between the two measurements. For MEG-MRI co-registration purposes, three anatomical landmarks (nasion, left pre-auricular, right pre-auricular), the locations of the marker coils, and 1000-5000 points from the head surface were acquired using a Polhemus Fastrak digitizer. A luminance-triggered photodetector output pulse was used to create a temporally precise timestamp upon the presentation of the visual grating.

### MEG Preprocessing

Data from two VSS patients and one control participant were contaminated by metal artefacts from non-removable dental implants or jewellery. Temporal signal space separation (0.9 correlation limit) was used to successfully suppress these artefacts in all cases.^40^ The remaining pre-processing was performed using the Fieldtrip toolbox v20191213.^41^ For each participant, the entire recording was bandpass filtered between 0.5-250Hz (Butterworth filter, 4^th^ order, applied bidirectionally) and band-stop filtered to remove residual 50Hz power-line contamination and its harmonics. Data were then epoched, based on the onset of the visual grating, into segments of 1.5s pre- and 1.5s post-stimulus onset. To avoid edge artefacts during time-frequency decomposition, an additional 2.5s of data on either side of these time-points was included as ‘padding’. MEG channels containing large amounts of artefactual data were identified by visual inspection (a maximum of ten channels, per participant, were removed).

Trials containing artefacts (SQUID jumps, eye-blinks, head movement) were removed by visual inspection. After pre-processing, there was an average of 109.7 trials (SD=9.1) for the VSS group and 117.4 trials for the control group (SD=2.3). Finally, data were down-sampled to 300Hz to speed computation.

### MEG-MRI Coregistration

As structural MRI scans were not available for all participants, we adopted an alternative approach for MEG-MRI co-registration. The digitised head-shape data were matched with a database of 95 structural MRIs from the human connectome database,^42^ using an iterative closest points (ICP) algorithm. The head shape-MRI pair with the lowest ICP error was then used as a ‘pseudo-MRI’ for subsequent steps. This procedure has been shown to improve source localisation performance in situations where a subject-specific anatomic MRI is not available.^43,44^

The aligned MRI-MEG image was used to create a forward model based on a single-shell description of the inner surface of the skull.^45^ In SPM12, a nonlinear spatial normalisation procedure was used to construct a volumetric grid (8mm resolution) registered to the canonical MNI brain.

### Source-Level Gamma and Alpha Power

Source analysis was conducted using a linearly constrained minimum variance beamformer,^46^ which applies a spatial filter to the MEG data at each point of the 8mm grid. Based on recommendations for optimising MEG beamforming,^47^ a regularisation parameter of lambda 5% was used. Beamformer weights were calculated by combining lead-field information with a sensor-level covariance matrix averaged across data from baseline and grating periods. Data were bandpass filtered between 40-70Hz (gamma) and 8-13Hz (alpha), and source analysis was performed separately. To capture induced rather than evoked visual power, a period of 0.3-1.5s following stimulus onset was compared with a 1.2s baseline period (1.5-0.3s before grating onset).

### ROI definition

To analyse changes in oscillatory power and PAC further, we defined a region of interest in the calcarine sulcus using the AAL atlas,^48^ which overlaps with visual area V1. This ROI was chosen based on previous MEG and intracranial recordings,^29,38,49,50^ which has established V1 as the primary cortical generator of gamma oscillations following the presentation of visual grating stimuli. For each participant, we selected the grid-point within the calcarine sulcus (parcel names: *Calcarine_L*; *Calcarine*_R), which showed the greatest change in gamma power versus baseline. The sensor-level data was then multiplied by the spatial filter from this grid-point to obtain a V1 “virtual electrode”.

### ROI Oscillatory Power and Peak Frequency

For the gamma band, oscillatory power was calculated using a multi-taper approach,^51^ from 40-70Hz, using a 0.5s time window, sliding in steps of 0.02s and ±7Hz frequency smoothing. For the alpha band, oscillatory power was calculated using a single Hanning taper between 8-13Hz, in steps of 1Hz, using a sliding window of 0.1s. The change in oscillatory power between baseline (−1.5 to −0.3s) and visual grating (0.3-1.5s) time-periods was averaged across 40-70Hz (gamma) and 8-13Hz (alpha) and expressed in decibels (dB). This time window was chosen to capture induced rather than evoked visual power. The frequency range 40-70Hz was chosen given previous research showing maximal changes in gamma oscillations for this frequency range.^29,39,50,51^ Post-hoc analysis across a wider frequency range (30-150Hz) confirmed that for our data, both groups showed maximal changes in gamma oscillations between 40-70Hz (see Supplementary Figure 1). To calculate the peak frequency of power changes for each participant, we used MATLAB’s *findpeaks.m* function between 40-70Hz (gamma) and 8-13Hz (alpha). Subject-specific results of this procedure are shown in Supplementary Figures 2a–b.

**Figure 2:**
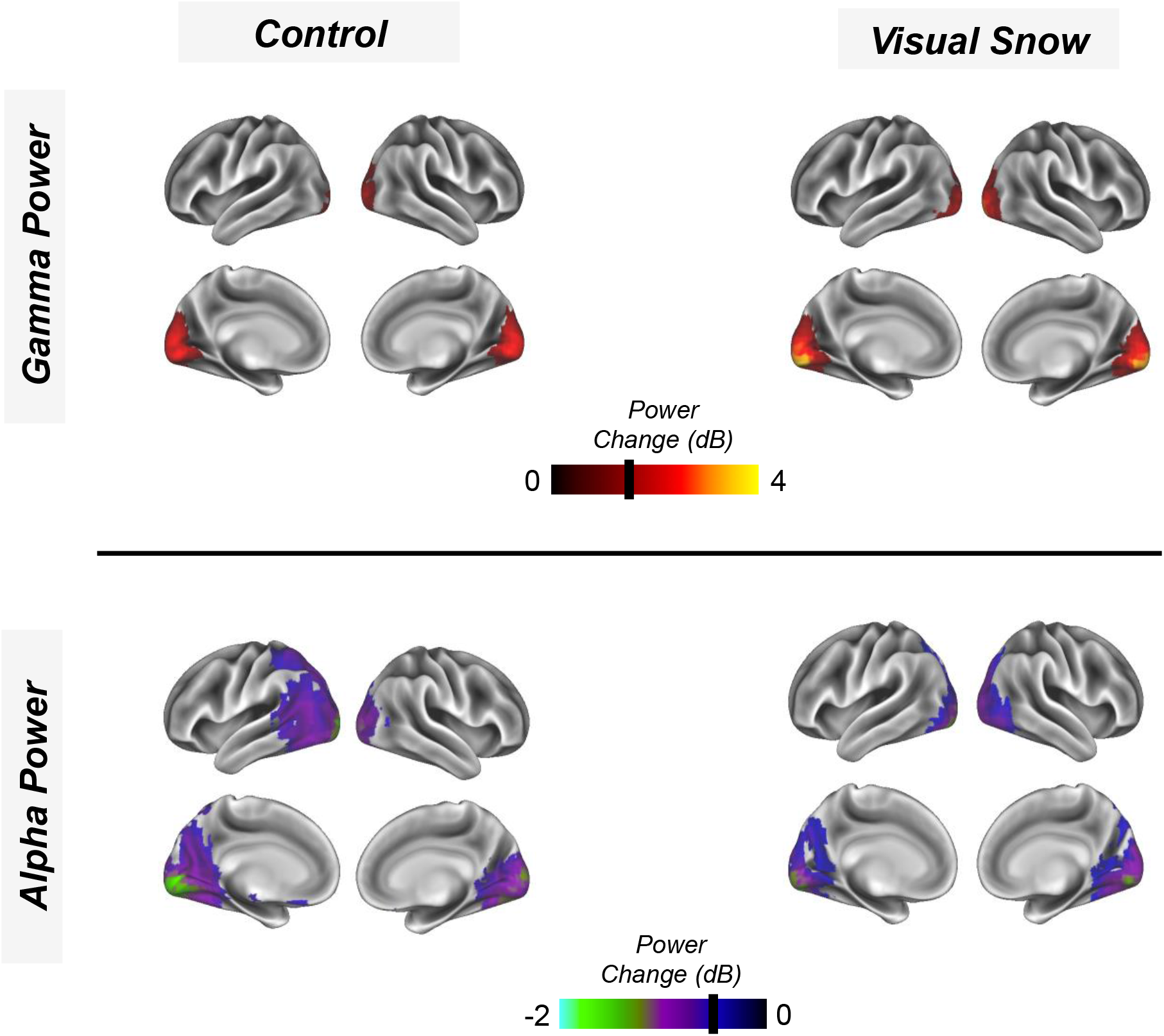
Following visual grating presentation, the change (dB) in gamma power (40-70Hz; 0.3-1.5s, upper panel) and alpha power (8-13Hz, 0.3-1.5s, lower panel) were calculated across a whole-brain grid. Results for the control group (left) and VSS group (right) were averaged and interpolated onto a 3D cortical mesh and finally thresholded at values greater than 1.3dB (gamma) and less than −0.3dB (alpha) for illustrative purposes.

### ROI Baseline Power

To check whether our results were driven by group differences in baseline power, for each subject, we averaged oscillatory power, as calculated in the previous section, between 1.5s to 0.3s before stimulus onset and 40-70Hz.

### V1 Phase-Amplitude Coupling (PAC)

Time courses from our ROI data were examined for changes in alpha-gamma phase-amplitude coupling (PAC). For a detailed discussion about PAC computation and methodological issues, see Seymour, Rippon, & Kessler (2017)^39^. Briefly, we calculated PAC values between phases 7-13Hz (in 1Hz steps) and amplitudes 34-100Hz (in 2Hz steps) for the time period 0.3-1.5s following the grating presentation. PAC values were corrected using 1.2s of data from the baseline period. This resulted in a 33*7 amplitude-phase comodulogram for VSS and control groups, which were statistically compared using a cluster-based permutation test.^52^ A more broadband frequency range for the amplitude was chosen so that we could capture the minimum and maximum edges of increased PAC in the comodulogram. To calculate PAC values, we used the mean vector length (MVL) approach from Ozkurt & Schnitzler^53^. Code used for PAC computation can be found at: https://github.com/neurofractal/PACmeg.

### Statistical Analysis

V1 oscillatory power and peak frequency were compared between groups using an independent samples t-test (two-tailed) implemented in JASP.^54^

For PAC, statistical analysis was performed using cluster-based permutation tests,^52^ which consist of two parts: first, an independent-samples t-test (two-tailed) is performed, and values exceeding an uncorrected 5% significance threshold are grouped into clusters. The maximum t-value within each cluster is carried forward. Second, a null distribution is obtained by randomising the participant label (VSS/control) 10,000 times and calculating the largest cluster-level t-value for each permutation. The maximum t-value within each original cluster is then compared against this distribution. The null hypothesis is rejected if the test statistic exceeds a threshold of p<.05 (corrected across both tails, i.e., p < 0.025 for each tail).

### Data availability

The data that support the findings of this study are available from corresponding author, CF, or first author, RS. Data can only be shared in a pre-processed and anonymised format, to comply with Macquarie University ethical guidelines.

## Results

### Epidemiology

The VSS cohort had a female-to-male ratio of 7/11 with ages ranging from 22 to 45 years old (mean age of 29 ± 7 years). Healthy controls consisted of 5 females and 11 males with ages ranging from 21 to 43 years old (mean age of 31 ± 6 years). The average symptom duration was 5 years for the VSS cohort, with 5 patients reporting symptoms since early teenage years. Associated visual and non-visual symptoms are summarised in Table 1. The VSS cohort consisted of 100% classic VSS with 94% reporting associated palinopsia, 61% photophobia, 72% nyctalopia, and 89% other positive visual phenomena. Associated non-visual symptoms included tinnitus in 94%, migraine in 61%, and tremor in 50% of patients. For the control group, 12.5% of the cohort reported symptoms consistent with migraine. No other clinical conditions were reported.

**Table 1:** Visual and non-visual symptoms reported by the VSS cohort.

| *Visual Symptoms* |  |
| --- | --- |
| Classic Visual Snow | 100% |
| Palinopsias | 94% |
| Photophobia | 61% |
| Nyctalopia | 72% |
| Positive Visual Phenomena | 89% |
| Duration of symptoms >1 year | 94% |

**Table 1:** Visual and non-visual symptoms reported by the VSS cohort.
| *Non-Visual Symptoms* |  |
| --- | --- |
| Tinnitus | 94% |
| Migraine | 61% |
| Tremor | 50% |

### Whole-Brain Alpha and Gamma Power

To demonstrate successful source localisation with our LCMV-beamformer pipeline,^46^ see *Materials and Methods*, we calculated changes in gamma power (40-70 Hz) and alpha power (8-13Hz), following presentation of the visual grating, across an MNI-warped whole-brain 8mm grid. Gamma power (40-70Hz) and alpha power (8-13Hz) were compared between 0.3-1.5s post-stimulus onset (to capture induced rather than evoked power) and a 1.2s baseline period. As expected, both the control and visual snow participants showed focal increases in gamma power (Figure 2, upper panel) for regions overlapping with primary visual cortex. Both groups also showed decreases in alpha power across the ventral occipital cortex (Figure 2, lower panel), consistent with previous studies.^50,51^

### V1 Gamma Power & Peak Frequency

A time-course from the grid-point showing the maximum change in gamma power within the calcarine sulcus (see *Materials and Methods*) was used for further analysis. An independent t-test was used to investigate group differences in gamma power (averaged across 0.3-1.2s, post-grating onset) and peak frequency. Results showed that gamma power was significantly greater in the VSS group (mean = 3.20dB) compared with the control group (mean = 2.27dB), t(32) = 2.147, p = .0395, d = .738 (also see Figure 3A). This result was not driven by differences in baseline gamma power between groups (see Supplementary Figure 3). There were no significant differences in gamma peak frequency between controls (mean = 52.63Hz) and VSS participants (mean = 53.17Hz), t(32) = 0.215, p = .831, d = .074 (also see Figure 3B).

**Figure 3:**
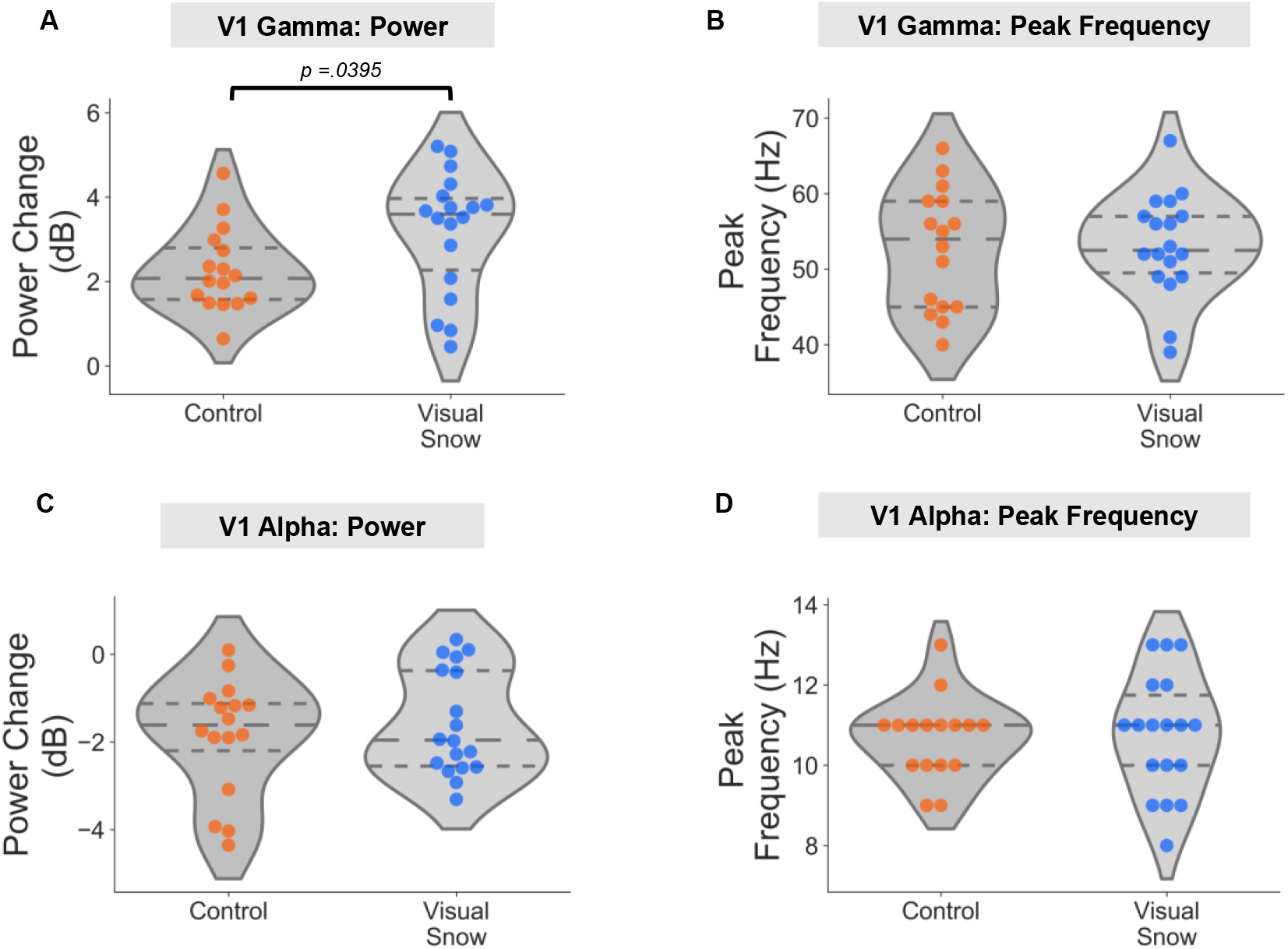
V1 Power and Peak Frequency. For both control and VSS groups, violin plots were produced (with median and interquartile range lines) to show: (A) V1 gamma power; V1 peak frequency; V1 alpha power; V1 alpha peak frequency. Dots correspond to data from individual participants.

### V1 Alpha Power and Peak Frequency

Using the same grid point, we repeated the analysis for the alpha band (8-13Hz), using an independent t-test to compare power and peak frequency between groups. There were no significant differences in alpha power between the VSS group (mean = −1.57dB) compared with the control group (mean = −1.99dB), t(32) = 0.873, p = .39, d = .30 (also see Figure 3C). There was also no significant difference in alpha peak frequency between groups (control mean = 10.7Hz; VSS mean = 10.8Hz), t(32) = 0.205, p = .84, d = .07 (also see Figure 3D).

### V1 Alpha-Gamma PAC

Using broadband data from V1, changes in alpha-gamma PAC were quantified using an amplitude-corrected mean-vector length algorithm,^53^ which has been shown to be robust for similar MEG data.^39,55^ For the control group, phase-amplitude comodulograms showed increased PAC following presentation of the grating versus baseline, peaking at 8–9Hz phase frequencies and 50–80 Hz amplitude frequencies (Figure 4, left). In contrast, the VSS group displayed lower changes in PAC across the comodulogram, with no clear positive peak (Figure 4, middle). Robust, non-parametric statistics were used to compare groups.^52^ For the control>VSS contrast, there was a single positive cluster of greater PAC between 8–9 Hz and 54–76 Hz, *p<.05 two-tailed* (Figure 4, right), i.e., coupling between alpha and gamma oscillations during perception in primary visual cortex is reduced in VSS compared to matched controls. We also quantified the effect size of this group difference, using Cohen’s d, see Supplementary Figure 4. The maximum value over the comodulogram was d = 1.24, which corresponds to a “very large” effect size.

**Figure 4:**
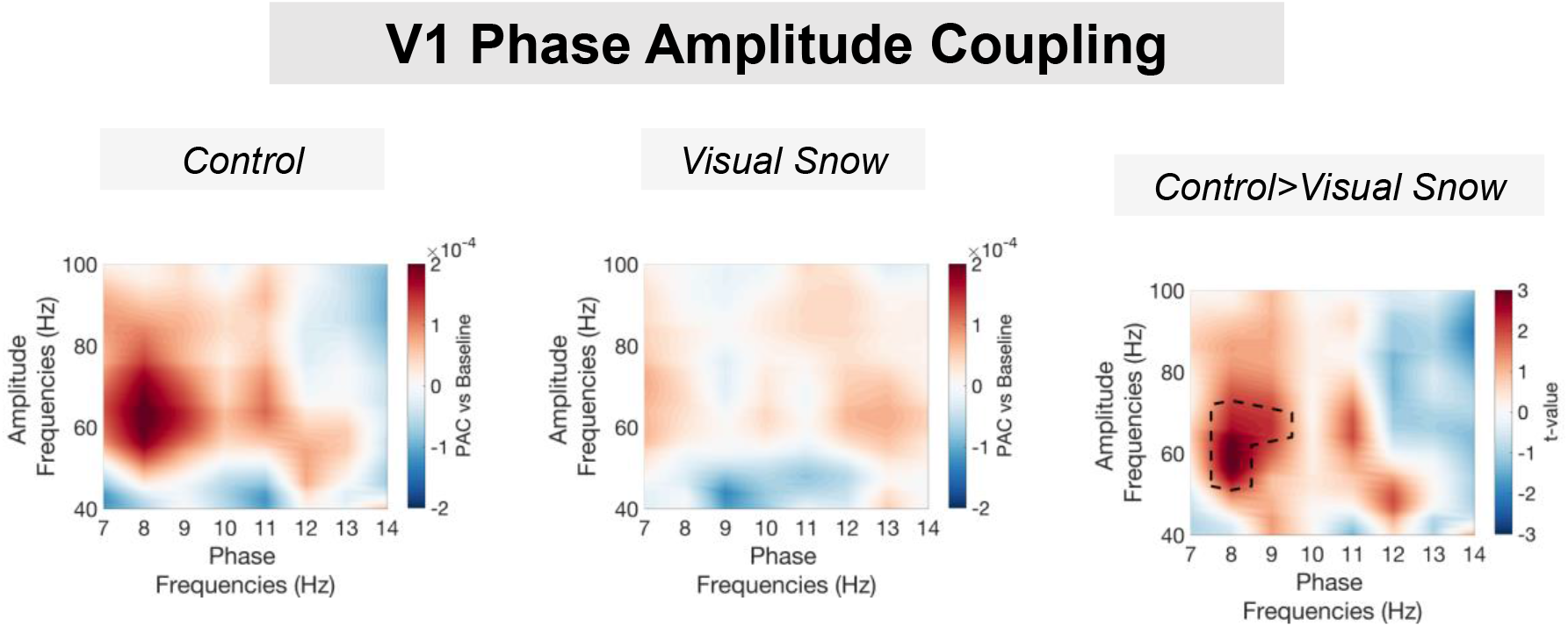
V1 phase-amplitude coupling. The control group showed increased alpha-gamma PAC compared with baseline, with a peak between 50–80 Hz amplitude and 8–9 Hz phase. The VSS group showed less prominent increases in PAC across the comodulogram. Non-parametric statistical comparison (see ‘Methods’) indicated significantly greater PAC for the control compared to the VSS group (p<.05) from 54–76 Hz amplitude and 8–9 Hz phase.

## Discussion

By utilising the excellent temporal resolution of magnetoencephalography, alongside beamforming for source localisation, this study supports our initial hypothesis that VSS may be considered a condition of visual dysrthymia.^1^

### Alpha-band (8-13Hz) oscillations in VSS

Occipital alpha rhythms dominate recordings made from resting healthy adults,^34^ and are involved in the active inhibition of irrelevant visual information.^35^ Reductions in alpha power measured using EEG/MEG are related to visual attention.^34,56^ Alpha is generally seen as an inhibitory rhythm; however, it is also linked with top-down modulation, prediction, and attentional sampling at ~10Hz.^20,57^ In this study, the presentation of a visual grating was accompanied by reductions in occipital alpha-band power, suggesting that participants were attending to the visual grating stimuli. However, there were no group differences in alpha power between VSS and control groups. We also investigated variation in individual alpha peak frequency, as peak alpha frequency is modulated by a variety of factors during perception.^58^ However, we found no differences in alpha peak frequency between groups.

Relating our findings to tinnitus, a related condition of phantom auditory perception, previous research has reported alterations in alpha-power and resting-state data.^31,59^ However, the literature is very heterogeneous, with both increases and decreases in alpha power being reported.^60–62^ Overall, it seems that neurophysiological mechanisms surrounding a ‘release from inhibition’ in the visual cortex (via alpha desynchronisation) are not directly involved in disorders of phantom perception. However, this does not rule out atypical mechanisms for top-down control via alpha-band *phase* relationships (see below: *Alpha-Gamma Phase Amplitude Coupling in VSS*).

### Gamma-band (40-70Hz) oscillations in VSS

Sensory stimuli elicit increases in high-frequency gamma oscillations generated through excitatory-inhibitory (E-I) neuronal coupling (see Buzsáki & Wang^27^) Gamma oscillations can be seen as a functional correlate of local neural ‘excitability’ and facilitate precise and effective inter-regional communication during sensory processing.^19,28^ Recent evidence suggests that gamma oscillations are primarily responsible for the feedforward flow of visual information up the cortical hierarchy.^63,64^

In this study, narrow-band (40-70Hz) oscillations originating from V1 were elicited using a high-contrast visual grating.^38,51^ We found that the VSS group had significantly greater gamma-band power compared to controls. The effect size of this finding was large: 13 out of the 18 VSS patients had gamma power values greater than the mean of the control group. Compared with controls, visual stimuli in VSS patients appear to elicit high-frequency, hyperexcitable activity in early visual cortex. We hypothesise that this hyperexcitable neural activity promotes atypical feedforward flow of information up the cortical hierarchy,^29,63,64^ manifesting as the disorganised white noise or ‘snow’ reported by VSS patients. These novel data highlight the advantages of studying VSS using MEG compared to EEG, where gamma oscillations are harder to measure.^38,50^

Alongside gamma power, we also calculated gamma peak frequency for each participant. Variability in gamma-peak frequency is determined by the balance between excitatory and inhibitory populations of neurons.^65^ However, we found no significant differences in gamma peak frequency between groups. Interestingly, there may be differential neural mechanisms behind the modulation of gamma amplitude versus frequency. Gamma peak frequency seems to be associated with the general “time-constant” of inhibitory processes in E-I circuits (Magazzini et al., 2016), whereas amplitude may be related to the strength of the inhibitory interneuron to superficial pyramidal cell connections.^66,67^

Our results generally complement those findings in a related and frequently co-existing condition: chronic tinnitus, where neuronal hyperexcitability and rapidly enhanced spontaneous firing rates are thought to result in excessive neuronal bursting and synchrony in the auditory cortex.^68,69^ This atypical neural synchrony is particularly linked with spontaneous gamma oscillations, commonly enhanced in tinnitus patients,^32,60,70^ and animal models of tinnitus.^71^ Increased sensory sensitivity, indexed via sensory-specific increases in gamma-band power, is a promising biomarker for disorders of phantom perception.

### Alpha-Gamma Phase Amplitude Coupling (PAC) in VSS

Emerging evidence has shown that the power (amplitude) of high-frequency cortical activity in primary sensory areas is modulated via the phase of lower-frequency oscillations.^72^ During visual processing, an increase in alpha-gamma phase-amplitude coupling (PAC) is frequently observed in electrophysiological recordings.^36,37^ Alpha-gamma PAC dynamically coordinates brain activity over multiple spatial scales,^73,74^ such that gamma oscillations within local neuronal ensembles are coupled with large-scale patterns of low-frequency phase synchrony.^75^ It is proposed that such dynamics allow information to be routed efficiently between brain areas and for neuronal representations to be segmented and maintained, e.g., during visual working memory.^76,77^

Following the presentation of a visual grating, we found that in VSS, alpha-gamma PAC in V1 was reduced compared to controls. This reduction occurred despite the VSS group displaying stronger visual gamma power in primary visual cortex. Interestingly, disruptions to PAC have also been reported in tinnitus,^78^ although increased PAC has also been shown.^79^

Our findings suggest that visual activity in VSS is both hyperexcitable (increased gamma power) and disorganised (reduced alpha-gamma PAC). Both results could be underpinned by an excitation-inhibition imbalance in visual cortex, as the neurophysiological generation of gamma amplitude and PAC relies heavily on local inhibitory populations of neurons.^80^ Affected local inhibitory processes would produce high-frequency ‘noisy’ activity and reduced signal-to-noise in perceptual systems, similar to findings reported in tinnitus.^16,81^ However, further corroborating evidence will be required before a definitive link between VSS, E-I interactions, and PAC can be confirmed. Disorganised local activity could also have concomitant effects on establishing inter-regional and global connectivity.^82^ Where top-down mechanisms are affected in VSS, altered noise-cancelling (i.e., the “gain”) of perceptual systems might result,^83,84^ meaning that typical visual stimuli would produce noisy and hyperactive responses in visual cortex, irrespective of their context.^1^ Reduced noise cancelling could explain previous EEG findings of reduced habituation in VSS.^14^ Future studies, specifically targeting perceptual gain and visual feedback pathways,^29,85^ should explore these ideas in more detail.

#### Clinical Relevance

From a clinical perspective, our novel findings of increased gamma power and reduced alpha-gamma PAC in VSS suggest that interventions targeting the re-establishment of typical rhythmical activity may help manage and treat the condition. Subject-specific neuromodulation approaches like repetitive TMS and cross-frequency transcranial alternating current stimulation,^86^ or neurofeedback approaches targeting gamma power and/or alpha-gamma PAC could be used for managing VS symptoms.^87,88^

#### Relation to other markers of VSS

Previous research has employed a range of imaging modalities to identify surrogate markers of brain dysfunction in VSS^89^. For example, using ^18^F-2-fluoro-2-deoxy-D-glucose PET, Schankin et al^8^ reported hypermetabolism in the lingual gyrus of VSS patients, alongside hypometabolism in the right superior temporal gyrus and the left inferior parietal lobule. Resting-state functional MRI data from a VSS cohort also showed hyperconnectivity between extrastriate and inferior temporal regions and between prefrontal and parietal cortex.^12^ It is tempting to link hypermetabolism and hyperconnectivity in VSS with our finding of increased gamma-band oscillations. However, the associations between visual gamma, BOLD, and PET are not well established. Generally, increased gamma power is related to increased BOLD ^90^, especially for broadband gamma responses ^91^. However, the relationship for narrow-band visual gamma is more nuanced (see: Muthukumaraswamy & Singh^50^; Singh^92^). It is also important to note that, unlike MEG, both PET and functional MRI data lack the temporal resolution required to measure dynamic changes to neural activity during visual perception.

Research utilising structural and functional MRI has reported disruptions to a wide array of brain regions in VSS. For example, increases in grey matter volume are found in lingual gyrus, fusiform gyrus junction, primary and secondary visual cortices, middle and superior temporal gyrus, and parahippocampal gyrus.^6,8,12^ Using functional MRI with a visual paradigm, Puledda and colleagues report decreased BOLD responses in VSS specific for the insula, which were interpreted as disruptions to the salience network.^6^ Overall, regions overlapping with extrastriate visual cortex seem to be most commonly associated with VSS.^6,8,12,89^ These regions are responsible for high-level visual processing such as colour vision perception and are linked with palinopsia^2^: a symptom that was present in 94% of our cohort. Our data extend this work by showing how functional changes in VSS are present even earlier in the visual hierarchy (i.e., primary visual cortex). These low-level alterations might then propagate downstream to extrastriate regions and beyond.

Finally, electrophysiological markers of VSS have reported a number of low-level differences versus controls, including increased N145 latency,^13^ and reduced habituation.^14,93^ Our results build on this research by demonstrating differences in the endogenous rhythms of the brain during visual processing. Findings of reduced habituation in VSS are particularly interesting, as they suggest a disrupted noise-cancellation mechanism, which is unable to modulate hyperactive and noisy V1 activity.

#### Thalamocortical dysrhythmia

While this study has focussed on dysrhythmias measured from the cortex, it is also essential to consider other brain regions, such as the thalamus. Work over the last few decades suggests that the thalamus does not simply act as a relay station during sensory processing. Instead, there exists a robust network of cortico-thalamic feedback neurons that dynamically influence sensory processing.^94^ One prominent theoretical account termed “thalamocortical dysrhythmia” (TCD) suggests that there is a final common pathway linking disorders of phantom perception, including e.g., migraine, tinnitus, neurogenic pain, and Parkinson’s disease,^25^ that slows the resting state alpha rhythm (8–13Hz) generated by the thalamus to just 4–7 Hz,^30^ and is accompanied by an increase in gamma power due to changes in lateral inhibition within thalamocortical circuits.^25,95^ In our cohort of VSS patients, we did not observe any slowing of alpha rhythms measured from the cortex; however, we did observe functionally increased gamma-band power, potentially related to changes in E-I interactions.^27,74,81^ Furthermore, our findings of reduced alpha-gamma PAC in VSS suggest that alpha-rhythms, typically generated by the thalamus, may become decoupled from gamma oscillations in the visual cortex.^25,37^ Interestingly, under the TCD framework,^25^ if thalamic rhythms have slowed to 4-7Hz in VSS, the visual cortex may become preferentially entrained to the theta rhythm (i.e., increased theta-gamma PAC). However, in this study, the length of each trial was insufficient to accurately quantify theta-gamma coupling.^39^

To further test the TCD framework, future work should focus on studying potential dysthymias directly within the thalamus and/or via thalamocortical connectivity. While, deep-brain structures like the thalamus are notoriously challenging to measure with non-invasive arrays of MEG sensors placed outside the head,^21^ recent progress has shown that it is possible,^96^ given certain constraints.^97,98^ However, in this study, the quality of the MEG-MRI co-registration, and the resulting forward model, were not good enough for reliably measuring subcortical activity. Therefore, future work should aim to utilise *subject-specific* 3D-printed scanner-casts and high-quality structural MRI scans in VSS cohorts.

### Limitations

Our study is based on a relatively small number of VSS and control participants. Participant recruitment was cut short by the COVID-19 pandemic. However, the effect sizes of group differences should be considered: d = .738 for gamma power (which can be described as “medium” to “large”); and d = 1.24 for the alpha-gamma PAC result (which can be described as “very large”). This strengthens our confidence in the inferences drawn from the results. Future studies should replicate and extend our findings, focusing on characterising dysfunctional oscillatory activity in VSS, with even greater precision. Larger cohorts of participants would also allow neuroimaging findings to be directly related to the clinical symptoms of the condition, a crucial consideration given that VSS exists on a continuum with significant variances in the severity of reported symptoms.^1,10^ Finally, this study opted to use a high-contrast visual grating to elicit specific visual oscillations in the early visual cortex. However, it remains unclear whether our findings generalise to more complex perceptual stimuli. Interestingly, VSS patients report that certain stimuli trigger “snow” symptoms more than others. More naturalistic stimuli (e.g., images and videos) combined with MEG could be used to isolate which particular aspects of the visual world intensify VSS symptoms. Immersive virtual reality environments could also be used in combination with new wearable MEG systems.^99^

## Conclusion

This study used MEG to study neuronal oscillations during visual processing in a cohort of visual snow syndrome (VSS) patients and control participants. Compared with controls, VSS patients displayed significantly increased gamma (40-70Hz) power in the primary visual cortex and reduced phase-amplitude coupling, suggesting that cortical activity in VSS during early visual processing is hyperactive and disorganised, results that are consistent with theories of thalamocortical dysrhythmia.

## Acknowledgements

We wish to thank all the patients and volunteers who gave their time to participate in this research study. We also acknowledge Nick Benikos and Stan Tarnavskii for MEG technical assistance.

## Supplementary material

This manuscript is accompanied by supplementary material.

## Supplementary Materials

**Supplementary Figure 1:**
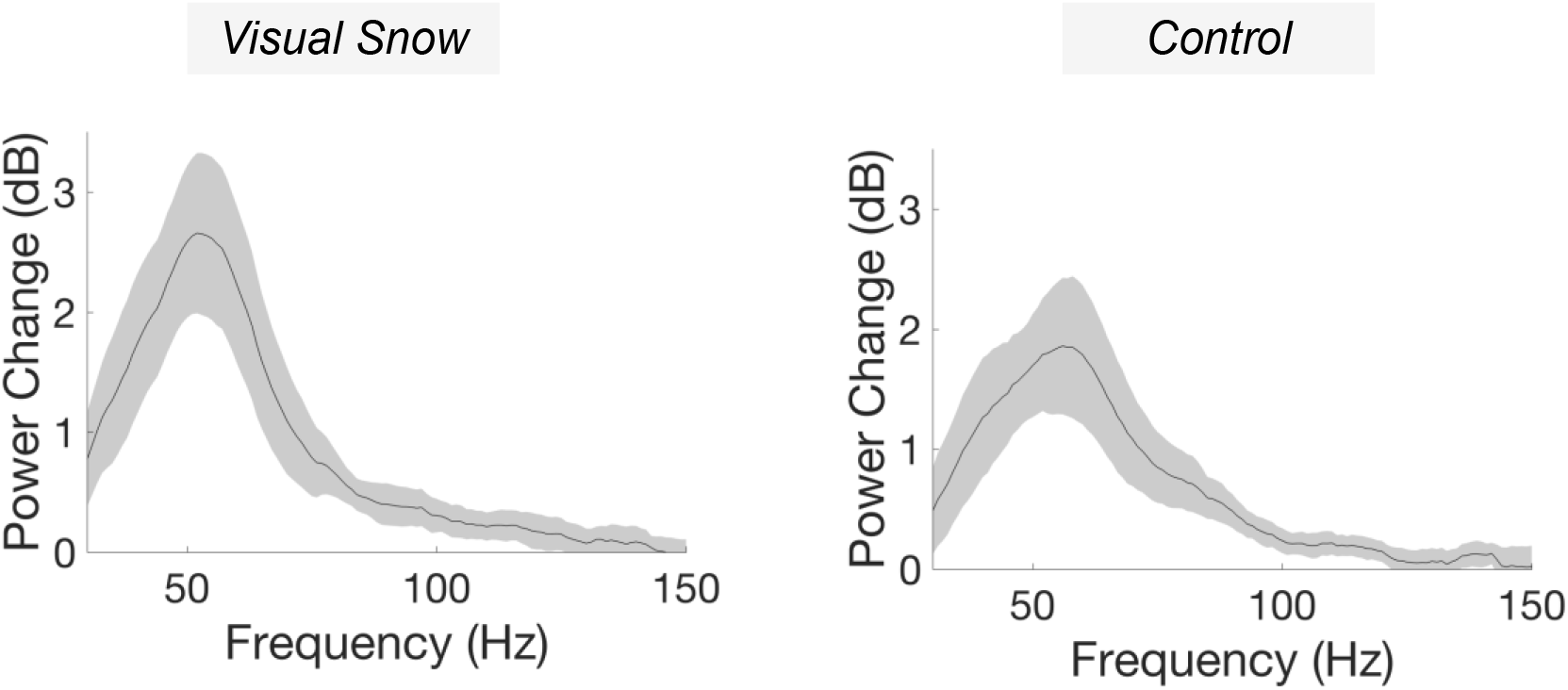
Gamma-band power change in the V1 region of interest, following presentation of the visual grating. The solid line represents the groups mean. Shaded error bars correspond to 95% confidence intervals. Note, the peak in the frequency spectrum from 40-70Hz across both groups.

**Supplementary Figure 2a:**
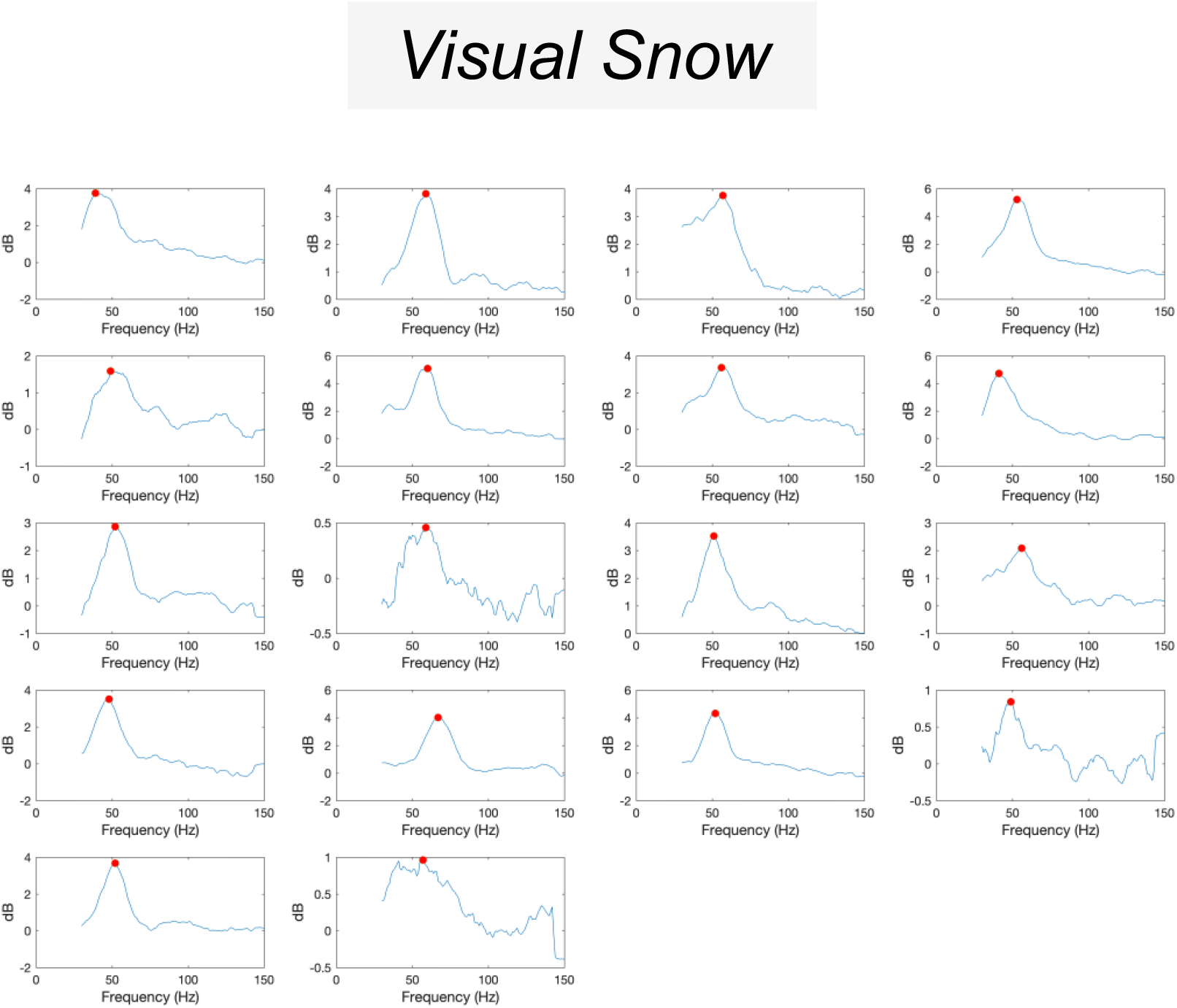
For each VSS participant, the change in gamma-band power following grating presentation in V1 is plotted, alongside the results of the peak-finding (using *findpeaks.m*)

**Supplementary Figure 2b:**
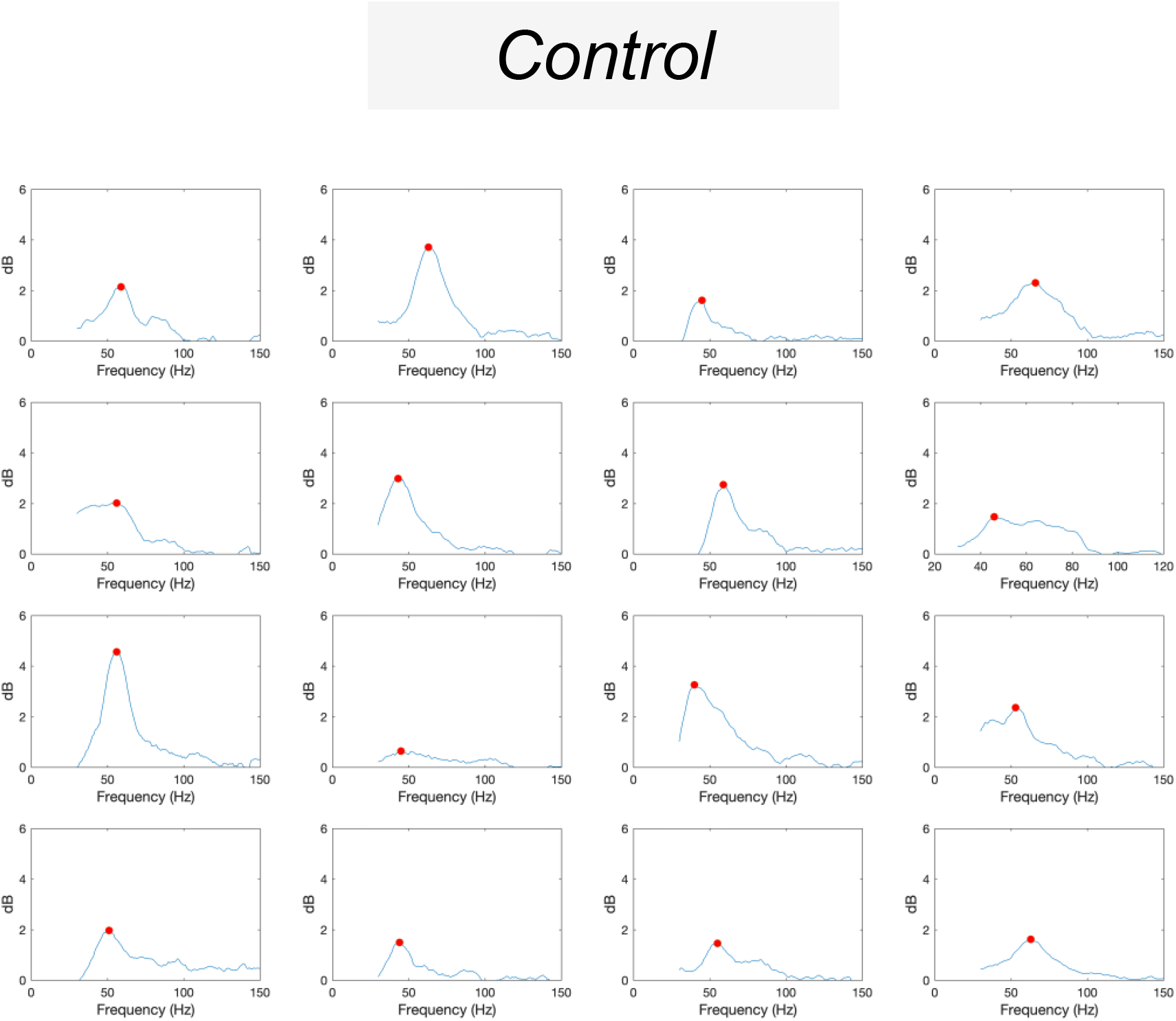
For each control participant, the change in gamma-band power following grating presentation in V1 is plotted, alongside the results of the peak-finding (using *findpeaks.m*)

**Supplementary Figure 3:**
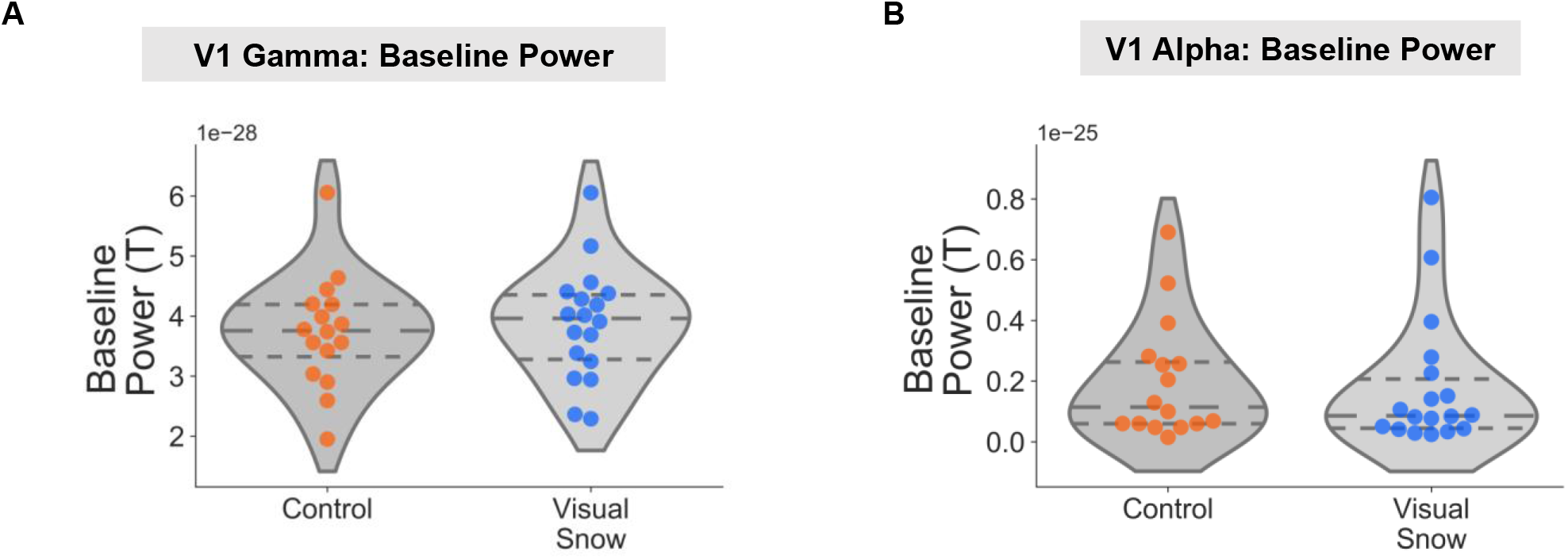
Data from the V1 region of interest was examined for potential group differences in baseline time-period (−1.5 to −0.2s relative to stimulus onset). No statistical differences in baseline power were observed between groups for (A) gamma or (B) alpha power, p>.05. Dots represent individual participants. Violin plots have the median and interquartile range shown with dotted lines.

**Supplementary Figure 4:**
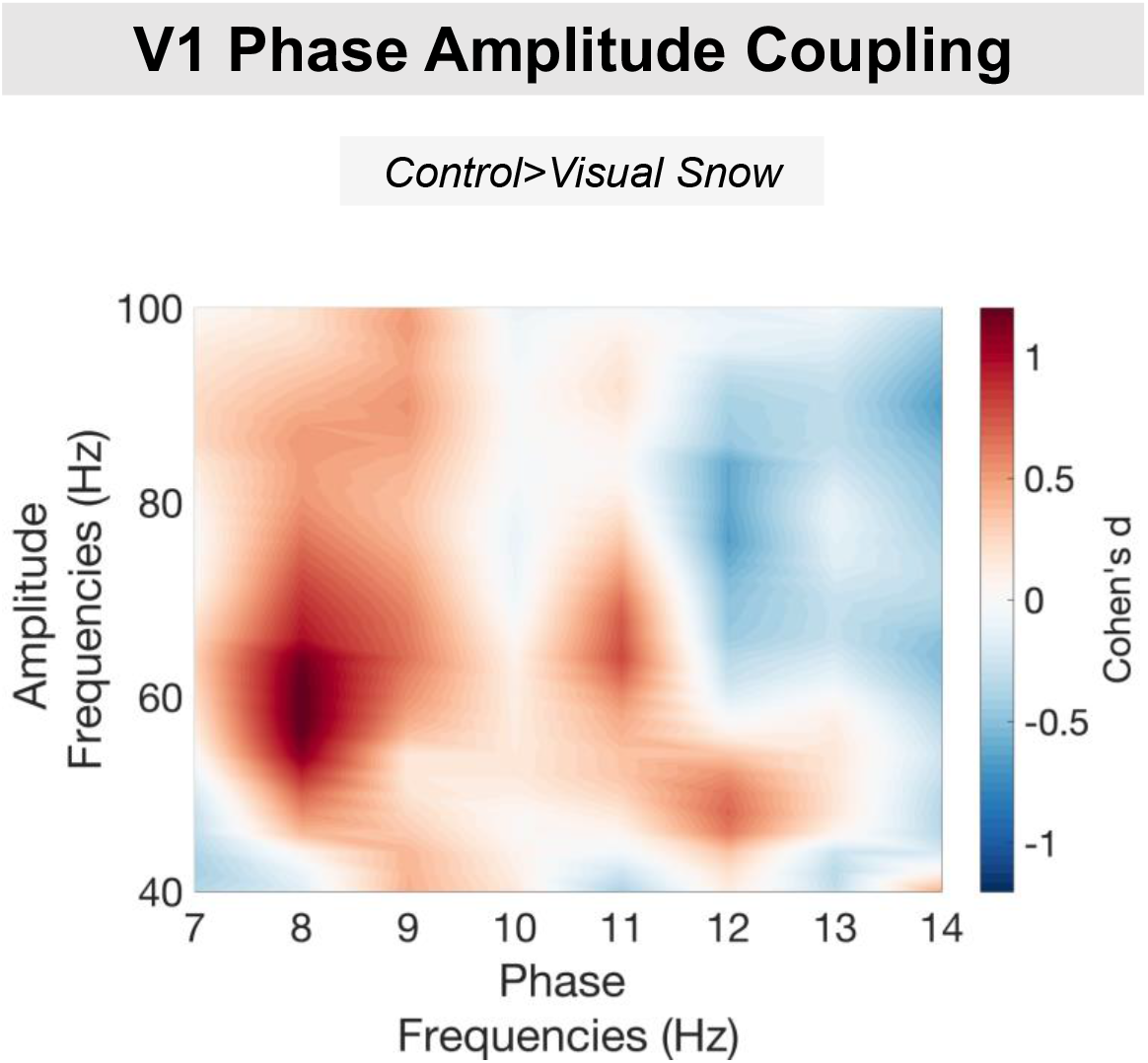
The effect size of the V1 PAC group difference was quantified using Cohen’s d (`ft_statfun_cohensd`). More details on the specific computational steps can be read here: https://www.fieldtriptoolbox.org/example/effectsize/

